## Supplemental Information for "Experience-dependent flexibility in a molecularly diverse central-to-peripheral auditory feedback system"

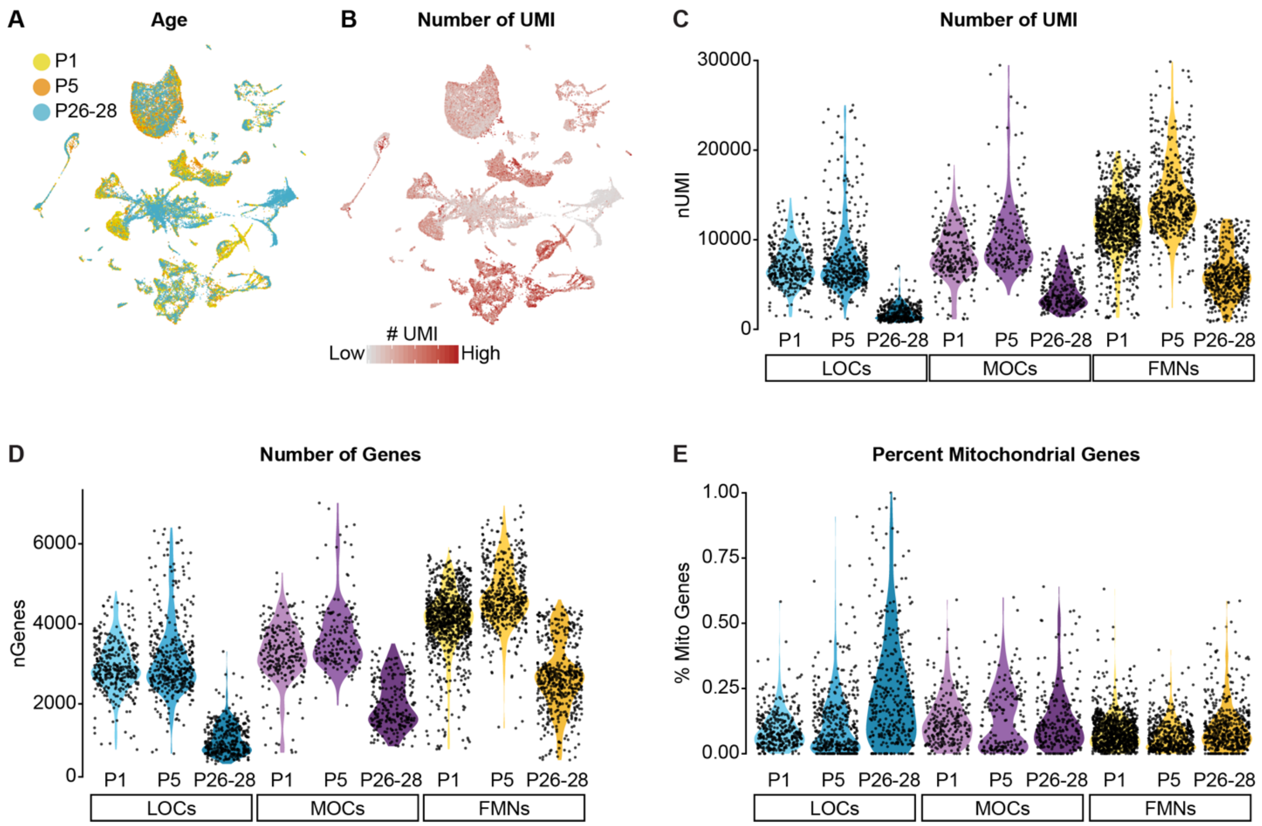


**Figure S1 (Related to Figure 1). Quality-control metrics for single-nucleus sequencing data.**
**(A)** UMAP plot showing contributions from each age collected. Each neuronal cluster includes cells from all three timepoints, indicating that the clusters are not being driven by variability across timepoints or batches.
**(B)** Feature plot detailing the number of unique molecular identifiers (UMI) detected in each cell. Although some cell types vary in their gene expression levels, the architecture of the data overall is not driven primarily by differences in gene expression or read depth.
**(C-E)** Violin plots denoting the number of UMIs (C) number of genes (D), and percentage of mitochondrial genes (E) detected per cell. Cells from P26-28 animals collected with 10x v2 kits consistently yielded a lower number of genes and UMI detected per cell compared to P1 and P5 datasets collected with 10x v3 kits.

### Figure S2 (Related to Figure 1)

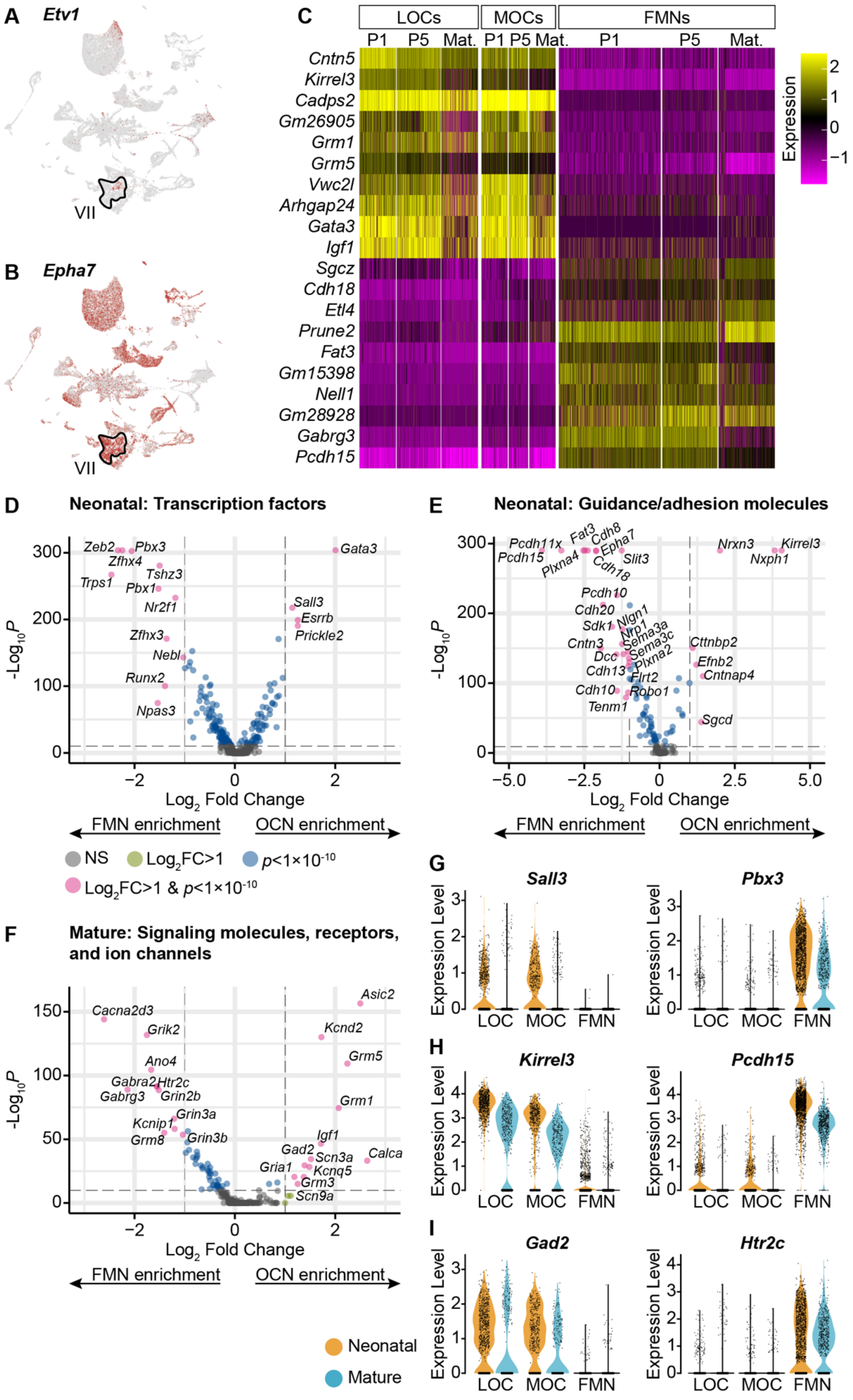

**Figure S2 (Related to Figure 1). OCNs and FMNs are transcriptionally distinct.**
**(A)** *Etv1* is a transcription factor selectively expressed in a subset of FMNs (VII). It is primarily expressed in only one motor neuron cluster.
**(B)** *Epha7* was previously identified as a marker for a subset of FMNs. Among the motor neurons in our dataset, it is expressed at high levels in the same cluster as *Etv1*, implying that cells in that cluster correspond to FMNs.
**(C)** Heatmap denoting the expression of 20 genes with significant differences in expression between OCNs and FMNs at both neonatal and mature timepoints ($p < 0.05$, Wilcoxon rank sum test, Bonferroni post-hoc correction). For visualization, scaled expression levels were capped at -2.5 and 2.5.
**(D-F)** Volcano plots denoting differential expression of transcription factors, guidance and adhesion molecules, and neuronal signaling components between OCNs and FMNs at P1-P5. Wilcoxon rank sum test, Bonferroni post-hoc correction.
**(G-I)** Violin plots denoting differentially expressed transcription factors (G), adhesion molecules (H), or signaling components (I) between OCNs and FMNs at neonatal (P1 and P5, orange) and mature (P26-28, blue) ages.

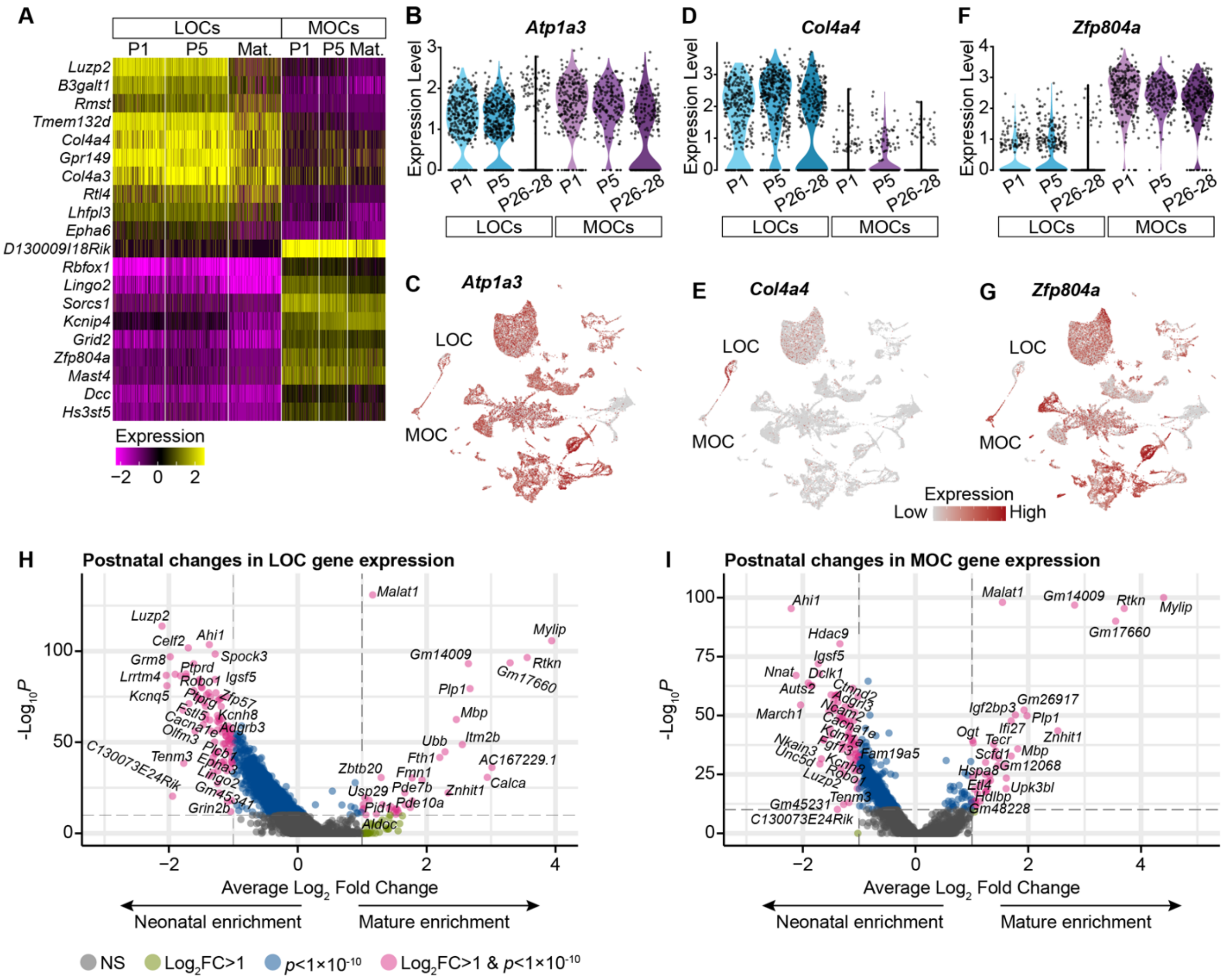


**Figure S3 (Related to Figure 2). OCN markers across postnatal development.**

**(A)** Selected genes with significantly different expression between MOCs and LOCs across postnatal development. ($p$ < 0.05, Wilcoxon rank sum test, Bonferroni post-hoc correction). For visualization purposes, scaled expression levels were capped at -2.5 and 2.5.

**(B, C)** Atp1a3 has been used as an MOC-specific marker in adult mice, but *Atp1a3* transcripts are broadly expressed in neonatal animals, including neonatal LOCs.

**(D, E)** *Col4a4* is enriched in LOCs across postnatal development.

**(F, G)** *Zfp804a* is expressed in multiple motor neuron subtypes, but in OCNs is preferentially expressed in MOCs at all postnatal timepoints.

**(H, I)** Volcano plots denoting gene-expression changes between neonatal (P1-P5) and mature (P26-P28) LOCs (H) and MOCs (I).

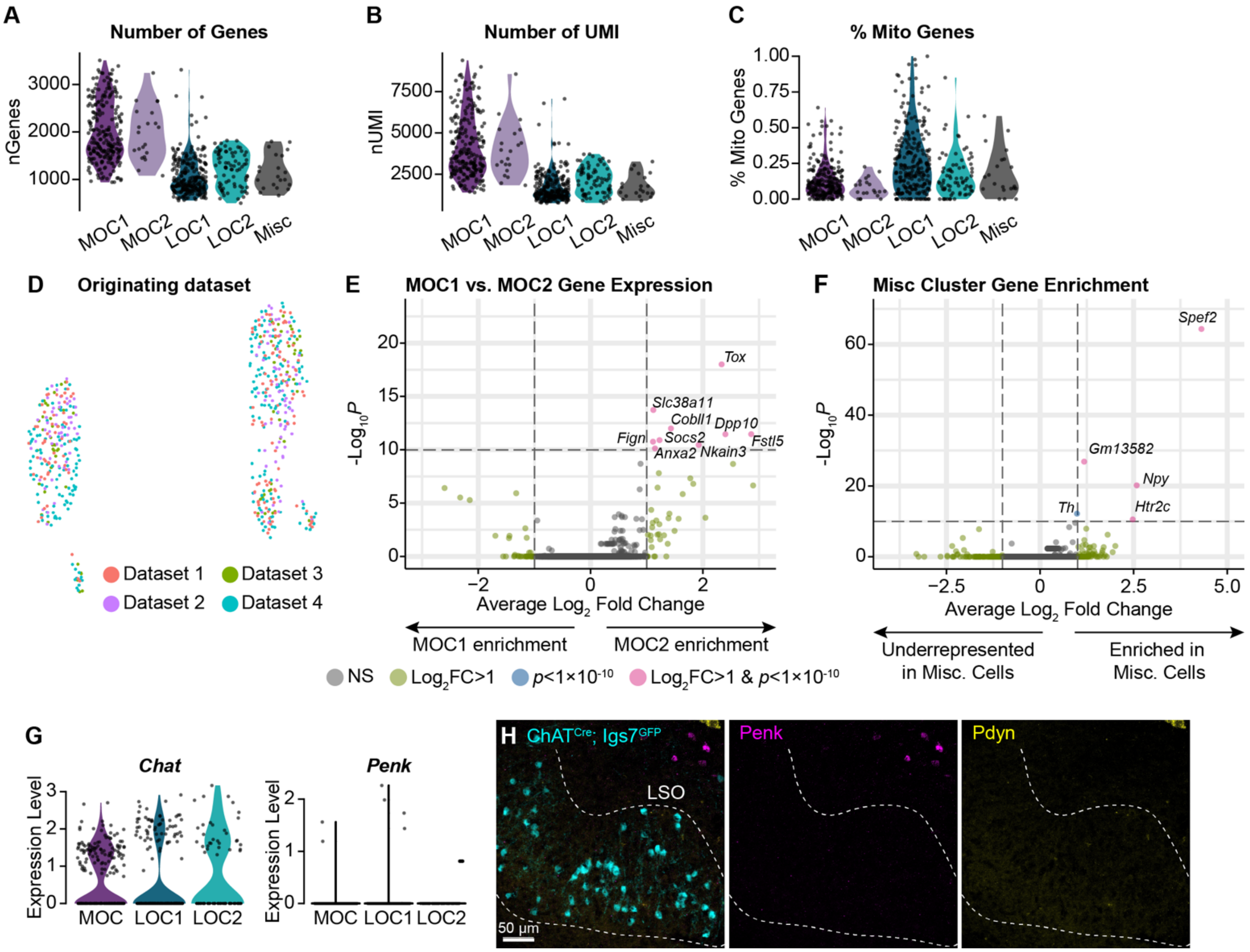


**Figure S4 (Related to Figure 3). Additional metrics for OCN transcriptional subtypes**.

**(A-C)** The number of genes detected (A), number of UMI (B), and percent of mitochondrial genes (C) is similar within MOC and LOC subtypes, indicating that differences between clusters are not reducible to technical variability.

**(D)** Cells from multiple collection rounds are not separable on a UMAP plot, suggesting that variability in OCN gene expression is not driven by batch-to-batch variation.

**(E)** Volcano plot summarizing gene-expression differences between MOC1 and MOC2 clusters.

**(F)** Volcano plot showing gene enrichment in the Misc cluster.

**(G, H)** Although our sequencing data detects similar levels of *Chat* in all OCN subtypes, we did not detect expression of *Penk* or *Pdyn* in either LOC subtype based on single-nucleus sequencing (G) or FISH (H). *Pdyn* did not pass our detection threshold for any adult OCN in our sequencing data, so no violin plot is shown. Non-LOC cells expressing *Penk* and *Pdyn* are visible in the upper right of (H). Representative images from 3 P28 $Chat^{Cre\Delta Neo}$; $Igs7^{GFP}$ mice.

**Figure S5 (Related to Figures 4 and 5)**

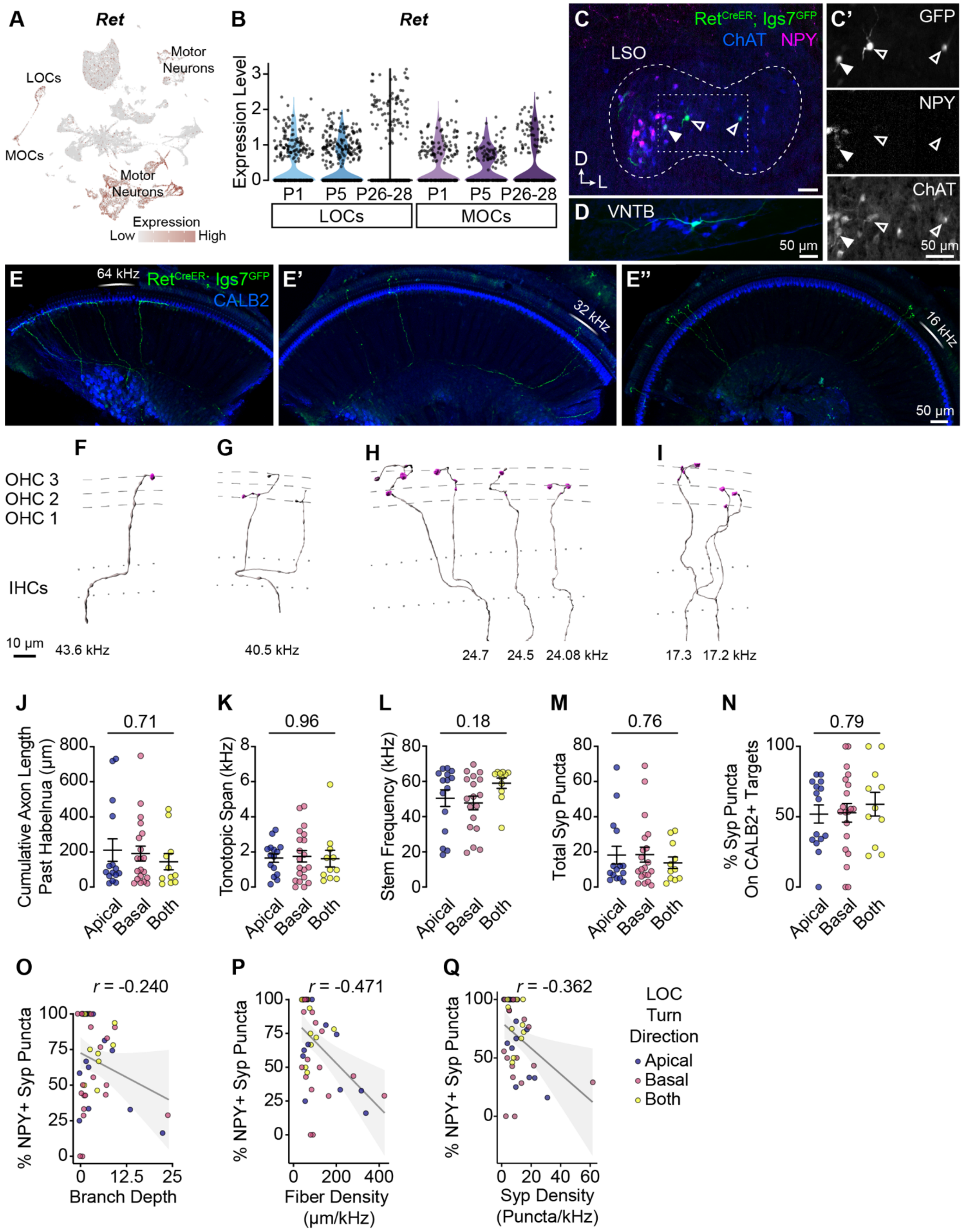

**Figure S5 (Related to Figures 4 and 5). Additional morphological quantification of OCN terminal axons.**

**(A, B)** *Ret* is broadly expressed in OCNs at least as early as P1. UMAP plot indicated *Ret* expression. *Ret* transcripts are detected in multiple motor neuron clusters, including OCNs (A). For visualization purposes, high expression on the color scale is capped at the $95^{th}$ percentile of gene expression. Violin plots indicate that *Ret* is expressed in OCNs across postnatal development (B).

**(C, D)** Representative sparse labeling of OCNs in the LSO (C) and VNTB (D) in $Ret^{CreER}$; $Igs7^{GFP}$ mice after a low dose of tamoxifen. Genetically labeled OCNs (green) colocalize with ChAT immunofluorescence (blue). LOC subtype is inferred based on NPY immunolabeling (magenta): both NPY+ (closed arrowhead) and NPY- (open arrowhead) LOCs are labeled (C').

**(E-E'')** Three segments from one cochlea (no OCN axons were present in the apex, not shown) with sparsely labeled LOC (E) and MOC (E'-E'') axons (green). CALB2 antibody (blue) labels SGNs and IHCs. This cochlea is from an animal with a single MOC neuron labeled in the brainstem.

**(F-I)** Imaris reconstructions of the MOC terminal axons from the cochlea in (E-E''), illustrating that a single MOC can have multiple branches innervating a ~27kHz frequency range from 43 to 17kHz. Dashed lines indicate OHC rows; dotted lines indicate the top and bottom of IHCs. Additional images of this same MOC are in Figure 4 A-A''. One axon from this neuron was bisected by the cut between cochlear turns and was not reconstructed.

**(J-N)** LOC axons that turn toward the base, apex, or both directions do not differ in total axon length (J), tonotopic span (K), stem frequency (L), number of Syp puncta (M), or Syp selectivity (the fraction of Syp puncta on CALB2+ targets, N). Error bars, mean ± SEM. Wilcoxon rank-sum test, n = 44 (K, L) or 46 (J, M, N) axons, N = 9 animals of either sex, P27-P30.

**(O-R)** There are no predictive relationships between the selectivity of LOC axons for CALB2+ targets and total number of Syp puncta ($r$ = -0.296, $p$ = 0.046) (O) or between the fraction of NPY+ Syp puncta and branch depth ($r$ = -0.240, $p$ = 0.108) (P). The fraction of NPY+ Syp puncta is weakly correlated with fiber density (cumulative μm of axon length/tonotopic span in kHz; $r$ = -0.471, $p$ = 0.001) (Q) and the density of Syp puncta ($r$ = -0.362, $p$ = 0.016) (R). $r$, Pearson's correlation coefficient. n = 46, N = 9 animals of either sex, P27-P30. Dot color corresponds to LOC axon turn direction.

**Figure S6 (Related to Figure 7)**

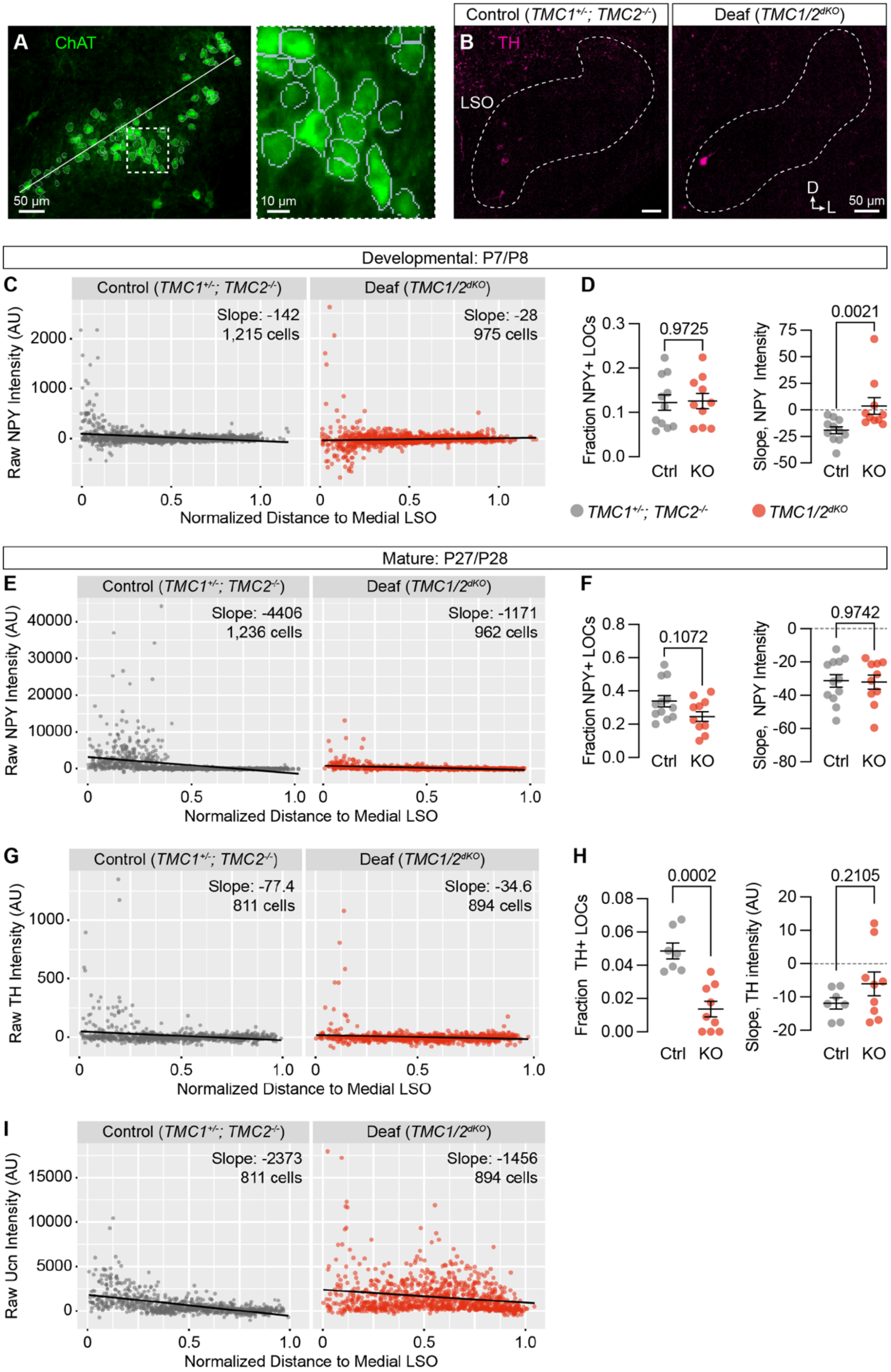

**Figure S6 (Related to Figure 7). Quantification of additional neurotransmitters in *TMC1/2$^{dKO}$* animals.**

**(A)** Example of the cell segmentation and size normalization procedure for quantifying expression of neurotransmitters in LOCs. White line indicates the length of the LSO used for distance normalization. Dashed line indicates region shown at higher power to the right, where gray boundaries indicate segmented cells.

**(B, C)** Representative images showing TH expression in the LSO of control (*TMC1$^{+/-}$; TMC2$^{-/-}$*) and deaf (*TMC1/2$^{dKO}$*) animals.

**(C-I)** Quantification of NPY (C-F), TH (G, H), and Ucn (I) fluorescence intensity, presented as a function of location within the LSO at developmental (C,D) or mature (E-I) stages. N = 7-12 LSOs from 5-6 animals per condition. Black line denotes regression line through all cells; slope denotes the slope of this regression line. Error bars, mean ± SEM; Wilcoxon rank-sum test.

**Video S1 (related to Figure 4). Terminal MOC axon morphology with putative synapses in the ISB and OHC region.** The full confocal z-stack volume of terminal MOC axons is shown, with CALB2 (blue) labeling IHCs and SGN peripheral fibers, the MOC axons labeled by *Ret*$^{CreER}$; *Igs7*$^{GFP}$ (white), and Syp labeling putative pre-synaptic contacts, which also delineate the three rows of OHCs (magenta). Using the GFP signal to create a mask of the Syp signal allows for reconstruction of the MOC axon and its associated putative pre-synaptic sites beneath the OHCs as well as in the ISB beneath the IHCs.

**Video S2 (related to Figure 5). Terminal LOC axon morphology with complex innervation patterns.** The full confocal z-stack volume is shown first, with CALB2 (blue) labeling IHCs and SGN peripheral fibers and the single LOC axon labeled by *Ret*$^{CreER}$; *Igs7*$^{GFP}$ (white), NPY (yellow), and Syp (magenta). The GFP signal is used to create a mask of the NPY and Syp channels. The axon is reconstructed and individual Syp puncta are categorized as On or Off a CALB2+ target, with Off-CALB2 Syp puncta generally localizing along the modiolar side of IHC row and On-CALB2 Syp puncta making contacts with the CALB2-high SGNs on the pillar side of the IHCs. Syp puncta are also coded as either NPY+ (lighter shade) or NPY- (more opaque).

**Table S1 (Related to Figure 3). Cell counts of NPY- and Ucn-expressing LOCs.** LOCs were manually annotated as expressing high or low levels of NPY or Ucn. Cells where both peptides were expressed above background are considered to be co-expressing both peptides. Each row includes cell counts from a single, 40 µm-thick section.

**Table S2 (Related to Figures 3, S4). Custom FISH probe sequences.** Oligo probe sequences used in Figure S4H for detecting *Penk* and *Pdyn* via hybridization chain reaction-based FISH.

| Measure | Pearson's *r* | *p* | $R^2$ | *n* (neurons) | Mean | SD |
|---|---|---|---|---|---|---|
| Membrane capacitance | -0.15 | 0.42 | 0.22 | 31 | 19.36 pF | 3.18 pF |
| Input resistance | 0.081 | 0.60 | 6.60E-03 | 45 | 324.6 MΩ | 186.02 MΩ |
| Outward K+ current magnitude | 0.1 | 0.53 | 0.01 | 43 | 2.78 nA | 1.62 nA |
| Steady-state inward current | -0.21 | 0.17 | 0.047 | 42 | -183.18 pA | 121.29 pA |
| A-type K+ current amplitude | -0.16 | 0.54 | 0.026 | 17 | 412.63 pA | 285.89 pA |
| Fast component of time constant of decay of fast-inactivating K+ current | -0.18 | 0.5 | 0.031 | 17 | 7.9 ms | 6.86 ms |
| Spontaneous firing rate | -0.28 | 0.35 | 0.08 | 13 | 8.6 Hz | 5.9 Hz |
| Rheobase | 0.06 | 0.76 | 0.004 | 24 | 13.3 pA | 8.20 pA |
| Action potential threshold | -0.25 | 0.26 | 0.06 | 24 | -49.08 mV | 6.12 mV |
| Amplitude from baseline of first spike evoked at rheobase | 0.01 | 0.95 | 0.0016 | 24 | 69.04 mV | 16.75 mV |
| Number of action potentials evoked by a 10 pA current injection | -0.08 | 0.75 | 0.006 | 24 | 4.5 | 3.11 |
| Input-output curve slope | -0.33 | 0.11 | 0.16 | 19 | 0.2 | 0.14 |

**Table S3 (Related to Figure 6). Physiological properties of LOCs do not correlate with CGRP-GFP fluorescence intensity.** Results of single linear regression and correlation analysis between CGRP-GFP and the indicated measures.

| Comparison | Calca | Calcb | Ucn | NPY |
|---|---|---|---|---|
| **LOC P1 vs. LOC P5** | <0.0001 | 0.1637 | 0.0728 | >0.9999 |
| **LOC P1 vs. LOC P28** | <0.0001 | <0.0001 | <0.0001 | <0.0001 |
| **LOC P1 vs. MOC P1** | >0.9999 | >0.9999 | >0.9999 | 0.5646 |
| **LOC P1 vs. MOC P5** | 0.0611 | 0.6236 | >0.9999 | >0.9999 |
| **LOC P1 vs. MOC P28** | <0.0001 | 0.0991 | >0.9999 | >0.9999 |
| **LOC P5 vs. LOC P28** | <0.0001 | 0.0202 | <0.0001 | <0.0001 |
| **LOC P5 vs. MOC P1** | <0.0001 | 0.0032 | 0.1595 | >0.9999 |
| **LOC P5 vs. MOC P5** | 0.0979 | >0.9999 | 0.7234 | >0.9999 |
| **LOC P5 vs. MOC P28** | 0.0755 | >0.9999 | 0.158 | >0.9999 |
| **LOC P28 vs. MOC P1** | <0.0001 | <0.0001 | <0.0001 | <0.0001 |
| **LOC P28 vs. MOC P5** | <0.0001 | 0.1575 | <0.0001 | <0.0001 |
| **LOC P28 vs. MOC P28** | <0.0001 | 0.3052 | <0.0001 | <0.0001 |
| **MOC P1 vs. MOC P5** | 0.1369 | 0.0332 | >0.9999 | >0.9999 |
| **MOC P1 vs. MOC P28** | <0.0001 | 0.0025 | >0.9999 | >0.9999 |
| **MOC P5 vs. MOC P28** | <0.0001 | >0.9999 | >0.9999 | >0.9999 |

**Table S4 (Related to Figure 7). Results of statistical tests comparing peptide expression between OCN groups at different ages.** Adjusted p-values from Kruskal-Wallis multiple-comparisons test for the indicated cell types and ages. Data is shown in Figure 7A-D.
